# Ryanodine Receptor Stabilization Therapy Suppresses Ca^2+^-Based Arrhythmias in a Novel Model of Metabolic HFpEF

**DOI:** 10.1101/2023.06.21.544411

**Authors:** Aaron D. Kaplan, Liron Boyman, Christopher W. Ward, W. Jonathan Lederer, Maura Greiser

**Affiliations:** Center for Biomedical Engineering and Technology, University of Maryland School of Medicine, Baltimore, Maryland, USA; Division of Cardiovascular Medicine, Department of Medicine, University of Maryland School of Medicine, Baltimore, Maryland, USA; Department of Physiology, University of Maryland School of Medicine, Baltimore, Maryland, USA; Greenebaum Comprehensive Cancer Center, University of Maryland School of Medicine, Baltimore, Maryland, USA; Department of Orthopedics, University of Maryland School of Medicine, Baltimore, Maryland, USA; Claude D. Pepper Older Americans Independence Center, University of Maryland School of Medicine, Baltimore, Maryland, USA

**Keywords:** Heart Failure, Arrhythmia, Calcium Handling, Metabolic Stress

## Abstract

Heart Failure with preserved ejection fraction (HFpEF) is the most prevalent form of heart failure worldwide and its significant mortality is associated with a high rate of sudden cardiac death (SCD; 30% - 40%). Chronic metabolic stress is an important driver of HFpEF, and clinical data show metabolic stress as a significant risk factor for ventricular arrhythmias in HFpEF patients. The mechanisms of SCD and ventricular arrhythmia in HFpEF remain critically understudied and empirical treatment is ineffective. To address this important knowledge gap, we developed a novel preclinical model of metabolic-stress induced HFpEF using Western diet (High fructose and fat) and hypertension induced by nitric oxide synthase inhibition (with L-NAME) in wildtype C57BL6/J mice. After 5 months, mice display all clinical characteristics of HFpEF and present with stress-induced sustained ventricular tachycardia (VT). Mechanistically, we found a novel pattern of arrhythmogenic intracellular Ca^2+^ handling that is distinct from the well-characterized changes pathognomonic for heart failure with reduced ejection fraction. In addition, we show that the transverse tubular system remains intact in HFpEF and that arrhythmogenic, intracellular Ca^2+^ mobilization becomes hyper-sensitive to ß- adrenergic activation. Finally, in proof-of-concept experiments we show *in vivo* that the clinically used intracellular calcium stabilizer dantrolene, which acts on the Ca^2+^ release channels of the sarcoplasmic reticulum (SR), the ryanodine receptors, acutely prevents stress-induced VT in HFpEF mice. Therapeutic control of SR Ca^2+^ leak may present a novel mechanistic treatment approach in metabolic HFpEF.

## Main Text

Heart failure (HF) with preserved ejection fraction (HFpEF), the most prevalent form of HF worldwide, is associated with significant morbidity and high mortality(1). Initially considered a disorder of diastolic dysfunction related to hypertensive left-ventricular remodeling, HFpEF is now understood as a heterogeneous disease widely seen in patients with significant metabolic stress driven by underlying comorbidities of obesity, dyslipidemia, hypertension and decreased insulin sensitivity (1, 2). A high rate of sudden cardiac death (SCD; ∼30-40 %) and ventricular arrhythmias contribute to HFpEF morbidity and mortality, and evidence that empiric anti-arrhythmic treatment is ineffective in HFpEF (3, 4) suggests that arrhythmias in metabolic stress-related HFpEF may arise via a distinct and as yet unidentified mechanism(3).

To investigate arrhythmia mechanisms pertaining to the large clinical cohort of metabolic stress related HFpEF we developed a novel preclinical model in wildtype mice (C57BL6/J). The model is based solely on mimicking the stressors (hypercaloric Western diet, hypertension, obesity) found in patients with metabolic stress related HFpEF, without relying on genetically altered animals. To achieve this, we added high dietary fructose (20% of dietary calories), an important driver of metabolic stress, to the well-established murine HFpEF model produced by high fat diet and hypertension-inducing nitric oxide synthase inhibition (with L-NAME)(5). After 5 months, this **f**ructose-metabolic HFpEF model (FM-HFpEF) demonstrated, *in vivo*, key aspects of the targeted clinical human HFpEF phenotype, including obesity, sarcopenia, decreased insulin sensitivity, high serum levels of brain natriuretic peptide (BNP), decreased global longitudinal strain (GLS) with normal ejection fraction (EF) and, importantly, stress-induced ventricular tachycardia (VT; Fig.1A-E).

**Figure 1.**
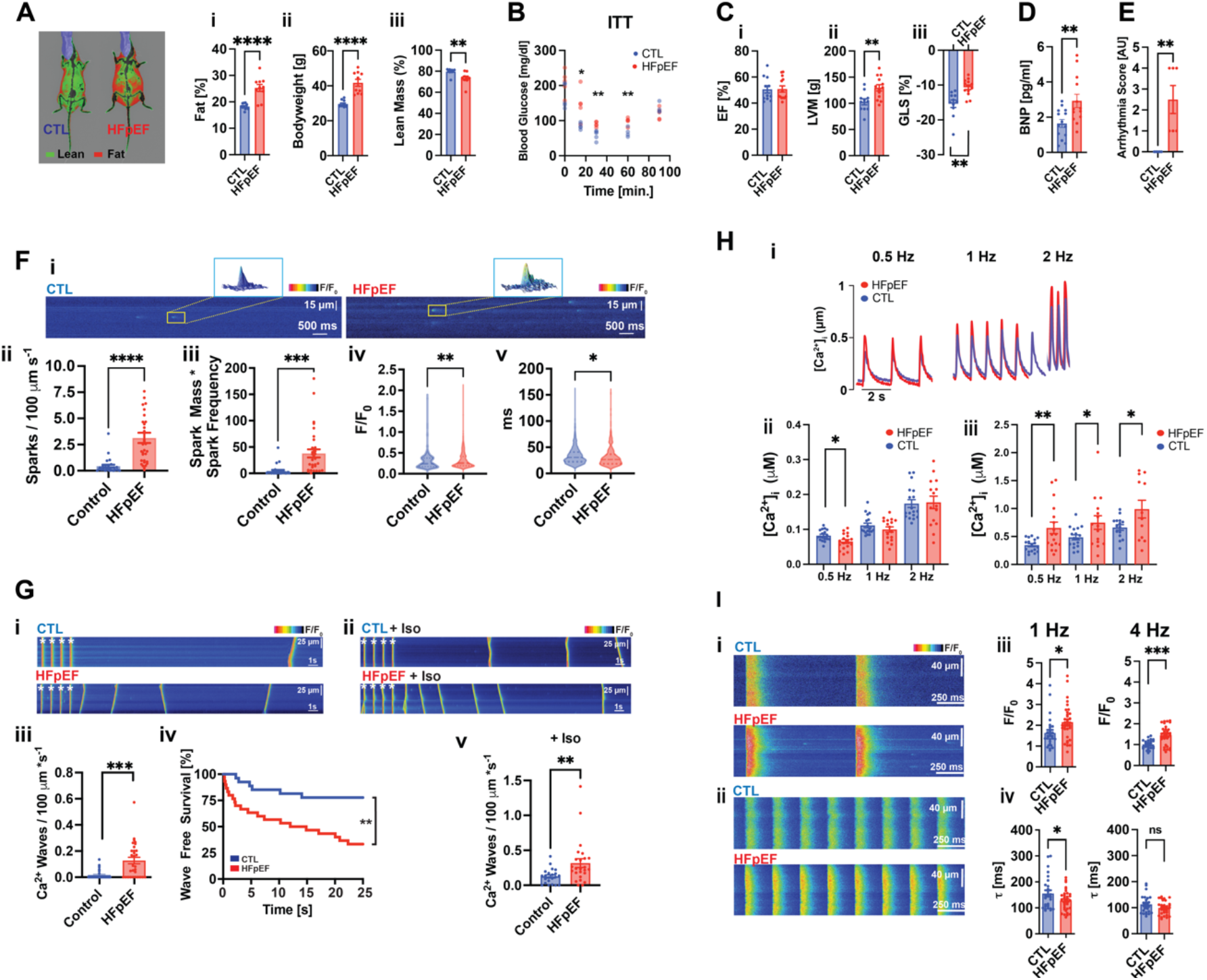
Arrhythmogenic Ca^2+^ signaling in metabolic HFpEF. Metabolic HFpEF model in mice. **A**. Body composition: **i**. Fat, **ii**. Bodyweight and **iii**. Lean Mass. **B**. Insulin tolerance test (ITT) **C**. Transthoracic echocardiography **i**. Ejection fraction (EF), **ii**. Left ventricular mass (LVM) **iii**. Global longitudinal strain (GLS), **: p<0.01. **D**. Serum levels of B-type natriuretic peptide (BNP). **E**. EKGs and arrhythmia score. **: p<0.01. Calcium imaging. **F**. Ca^2+^ sparks in HFpEF. **i**. Exemplars of Ca^2+^ sparks. **ii**. Ca^2+^ spark frequency, **iii**. Ca^2+^ spark-mediated SR Ca^2+^ leak, **iv**. Ca^2+^ spark amplitude, **v**. Ca^2+^ spark duration as full duration half maximum (FDHM). ****: p<0.0001; ***: p<0.001; **: p<0.01; *: p<0.05. **G**. Ca^2+^ waves. **i**. Exemplars of spontaneous Ca^2+^ waves. **ii**. Exemplars of spontaneous Ca^2+^ waves after isoproterenol treatment. *denotes stimulated beat. **iii**. Ca^2+^ wave frequency **iv**. Ca^2+^ wave free survival **v**. Ca^2+^ wave frequency (+ isoproterenol). ***: p<0.001; **: p<0.01. **H**. [Ca^2+^]_i_ in HFpEF myocytes. **i**. Stimulated Ca^2+^ transients (0.5-2 Hz). Summary data of **ii**. diastolic [Ca^2+^]_i_, **iii**. Systolic [Ca^2+^]_i_. **I. i, ii**. Stimulated Ca^2+^ transients (1-4 Hz). **iii**. Ca^2+^ transient amplitude **iv**. Ca^2+^ transient decay.

Ca^2+^ “leaking” from its stores in the sarcoplasmic reticulum (SR) is the key mechanism underlying Ca^2+^ based arrhythmogenesis in heart failure with reduced ejection fraction (HFrEF)(6, 7). Here we show a massive arrhythmogenic intracellular Ca^2+^ leak in FM-HFpEF ventricular myocytes, much larger than in HFrEF, evidenced by a ∼ 10-fold increase in spontaneous Ca^2+^ spark frequency and ∼ 9-fold increase in spark mediated SR Ca^2+^ leak (Fig.1F). Consistent with this massive SR Ca^2+^ leak was a dramatic increase in spontaneous Ca^2+^ wave activity (∼7-fold higher in FM-HFpEF myocytes compared to control myocytes, Fig.1G), the cellular trigger for Ca^2+^ based arrhythmias(6, 7).

We show that ß-adrenergic challenge *in vivo* elicits VT in FM-HFpEF mice (Fig.1E). We also find that *in vitro* ß-adrenergic challenge (isoproterenol 50 nmol/L) in single FM-HFpEF cardiomyocytes elicited a much higher frequency of arrhythmogenic Ca^2+^ waves than in control cells, thus further supporting the Ca^2+^ basis of arrhythmias in metabolic HFpEF (Fig.1G). Importantly, this high degree of intracellular Ca^2+^ mobilization *in vitro*, together with the inducibility of sustained VTs *in vivo*, demonstrate an intact and highly sensitive ß-adrenergic response in FM-HFpEF.

A crucial underlying mechanism of Ca^2+^ based arrhythmogenesis in HFrEF is a higher diastolic Ca^2+^ concentration in the cytosol ([Ca^2+^]_i_), which sensitizes the Ca^2+^ release channels of the SR, the ryanodine receptors type 2 (RyR2) to release Ca^2+^ (8). Surprisingly, calibrated measurements of [Ca^2+^]_i_ in single cardiomyocytes electrically paced at physiological frequencies showed that diastolic [Ca^2+^]_i_ was not increased in FM-HFpEF myocytes (2 Hz, Fig.1Hi,ii) despite an increase in the magnitude of their systolic [Ca^2+^]_i_ (Fig.1Hi-iii). In addition, Ca^2+^ transients from FM-HFpEF mice were shorter or unchanged from controls at physiological stimulation rates of 1-4 Hz, another striking difference from HFrEF where Ca^2+^ transients are prolonged (Fig.1I). To define the underlying mechanisms of this FM-HFpEF Ca^2+^ phenotype, we first examined the ultrastructure of the transverse and axial tubular system (TATS) of living, single cardiomyocytes. Sub-diffraction Airy scan imaging revealed no changes in TATS density, organization or orientation in FM-HFpEF (Fig.2A). These results are contrary to the severe reduction and disorganization of TATS pathognomonic for HFrEF, where the structural disruption of the cardiac dyad, the close apposition (∼20 nm) of L-type Ca^2+^ channels (Ca_v_1.2) with RyR2, causes arrhythmogenic SR Ca^2+^ release (9).

We next quantified the expression and post-translational modification (PTM) of dyadic and SR Ca^2+^ handling proteins. In ventricles from FM-HFpEF mice Ca_v_1.2 expression was unchanged. Surprisingly, RyR2 expression and its phosphorylation (Serine2808) were significantly reduced (Fig.2B), another important distinction from changes seen in HFrEF. In addition, calsequestrin 2 (CSQ2), the major intra-SR Ca^2+^ buffering protein, was significantly reduced, as was the expression and phosphorylation of phospholamban (PLB), the regulatory protein of the SR Ca^2+^ ATPase (SERCA) (Fig.2B). Taken together, these results suggest a significant remodeling of the cardiac dyad due to an altered stoichiometry of the Ca_v_1.2-RyR2 nanodomain. In addition, the expression and PTM of key SR Ca^2+^ handling proteins are reduced (schematic shown in Fig. 2D, E). While future work is necessary to decode the ensemble contribution of these changes to the distinct HFpEF Ca^2+^ signaling, reduced CSQ2 is known to cause ventricular arrhythmias via SR Ca^2+^ leak in cardiomyocytes due to reduced intra SR Ca^2+^ buffering (10).

**Figure 2.**
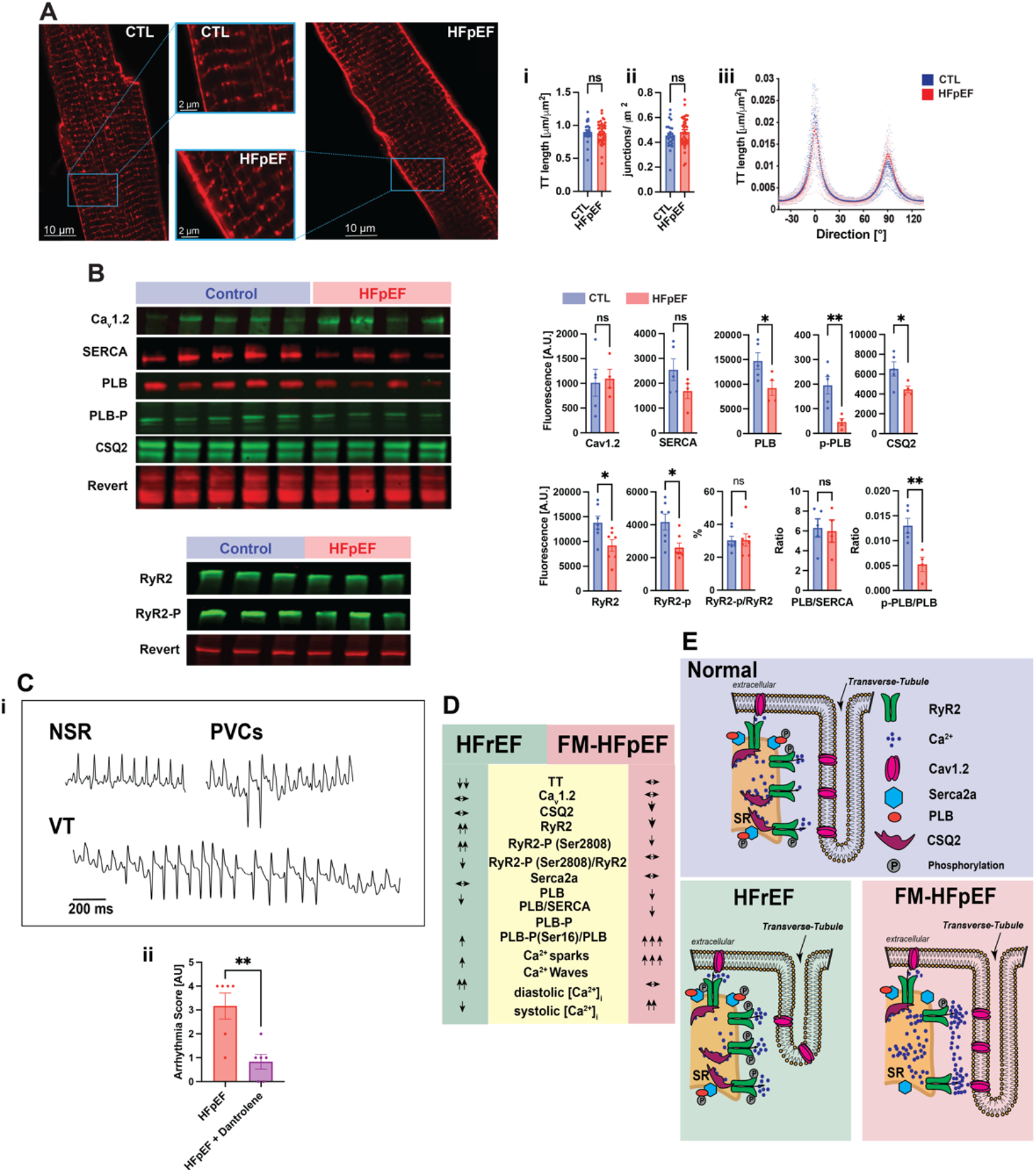
**A**. Transverse-Axial Tubule system (TATS). AiryScan images of living ventricular myocytes stained with the membrane dye di-8-ANEPPS (5 μmol/L) to label the TATS. **ii**. Total length of the TATS. **iii**. Number of junctions that connect intersecting perpendicular tubules. **iv**. The length of the TATS at each direction within the cells. **B**. Protein expression levels of key Ca^2+^ handling proteins. Ca_V_1.2: α1c subunit of the L-type Ca^2+^ channel, SERCA: SR Ca^2+^ ATPase, PLB: phospholamban, PLB-P: phosphorylated-PLM (Serine16), CSQ2: calsequestrin 2, RyR2: Ryanodine receptor type 2, RyR2-P: phosphorylated RyR2 (Serine2808). **C**. *in vivo* dantrolene treatment of FM-HFpEF mice. **i**. Summary data showing arrhythmia index in FM-HFpEF mice (n=6) pretreated with i.p. dantrolene (40 mg/kg i.p; 30 min) compared to untreated FM-HFpEF mice (n=6). **L**. Table showing changes in intracellular Ca^2+^ handling in HFrEF compared to the FM-HFpEF model. **M**. Schematic depicting organization of the dyadic Ca^2+^ release-apparatus in normal, HFrEF and FM-HFpEF.

To test a novel mechanistic approach for anti-arrhythmic therapy in HFpEF we performed proof-of-concept experiments with dantrolene, a ryanodine receptor stabilizing agent. Dantrolene is used clinically for the treatment of malignant hyperthermia (11) and has recently been shown to suppress Ca^2+^ based arrhythmias in human hearts *ex vivo* (12). Here we show that acute, mechanistic Ca^2+^ stabilizing treatment with dantrolene of FM-HFpEF mice *in vivo* (40 mg/kg i.p) significantly reduced arrhythmia burden and completely abolished stress-induced VTs (Fig.2C).

In summary, we report a novel preclinical model of metabolic HFpEF that phenocopies the complex cardiometabolic features and stress-induced ventricular tachycardias seen in clinical HFpEF related to metabolic stress (Fig.1A-E) (1, 3). We show here for the first time that metabolic stress induced HFpEF produces Ca^2+^ based arrhythmogenesis distinct from the “Ca^2+^ phenotype” pathognomonic for HFrEF (Fig.2D, E). Specifically, a massive intracellular Ca^2+^ leak, unchanged diastolic [Ca^2+^]_i_, intact TATS and reduced expression of key SR Ca^2+^ handling proteins underlie arrhythmogenic Ca^2+^ signaling in HFpEF. Importantly, these changes are also distinct from a form of HFpEF secondary to genetic hypertension (Dahl salt-sensitive rats) that progresses to HFrEF, where the ß-adrenergic responsiveness of intracellular Ca^2+^ mobilization was blunted (13). Another novel and distinct (from HFrEF) finding in FM-HFpEF was a decrease in CSQ2 expression. Previous work established that even small reductions of CSQ2 expression produce significant Ca^2+^ dependent arrhythmias, due to both the reduced Ca^2+^ binding capacity of the SR and increased RyR2 activity (10).

Finally, we demonstrate in proof-of-concept experiments that dantrolene, a clinically relevant ryanodine receptor stabilizer, acutely abolishes stress-induced VTs in FM-HFpEF mice. These results represent the first mechanistic, antiarrhythmic treatment concept (RyR2 stabilization) for VTs in metabolic HFpEF. Importantly, dantrolene, which is contraindicated in HFrEF due to its negative inotropy, could be used clinically in HFpEF patients. Future work will further evaluate the long-term effects and clinical treatment potential of ryanodine receptor stabilization therapy in metabolic stress related HFpEF.

## Materials and Methods

Male C57BL6/J mice were used in this study. Cardiac myocyte isolation, Ca^2+^ imaging and sub-diffraction microscopy were performed as previously described(14, 15). Detailed descriptions are provided in the Supporting Information.

## Acknowledgments

Sources of Funding: University of Maryland Claude D. Pepper Center Grant P30 AG028747 (MG).7U19 AI090959 (WJL &LB); Frontiers in Anesthesia Research Award from International Anesthesia Research Society (WJL & LB); R01 HL142290 (WJL &CWW); 5R35GM140822 (WJL); U01 HL116321 (WJL); Special BioMET/UMB Funds (WJL). R01 AR071618 (CWW).

