## Supplemental Material for "Ryanodine Receptor Stabilization Therapy Suppresses Ca^2+^-Based Arrhythmias in a Novel Model of Metabolic HFpEF"

**This PDF file includes:**

Supporting text (Extended methods)
SI References

**Methods**
All experiments involving animals were approved by the Institutional Animal Care and Use Committee of the University of Maryland School of Medicine, Baltimore, Maryland. Statistical analysis was performed using GraphPad Prism v9 and Matlab. Results are presented as mean ± SEM. No statistical analyses were performed to predetermine sample sizes; estimates were made on the basis of our previous experience, experimental approach, and availability and feasibility required to obtain statistically significant results. When representative images are shown, the selected images are those that most accurately represent the average data obtained in all samples.

**Body composition**
Body composition was quantified *in vivo* with an iNSIGHT DXA (Scintica, Canada) using dual x-ray absorptiometry.

**Insulin tolerance test (ITT)**
ITT was performed (0.5 U/kg recombinant human insulin I.P) after a 4 hour fast with glucose monitored (Contour Next One; time 0, 15, 30, 60 and 90min) from tail blood in HFpEF (n=5) and control (n=5) mice.

**Echocardiography**
Transthoracic echocardiography was performed as described previously using a VisualSonics Vevo 2100(1). Mice were anesthetized using inhalation anesthesia with isoflurane. Body temperature was monitored throughout acquisition. Systolic parameters were obtained from short-axis M-mode scans at the midventricular level, as verified by papillary muscles. Apical four-chamber views were obtained, and diastolic function was assessed by pulsed wave doppler imaging across the mitral valve. B-mode imaging in the parasternal long axis plane was used to calculate global longitudinal strain with VevoStrain software (Visual Sonics). At the end of the acquisition all animals recovered without issues.

**Brain natriuretic peptide (BNP) measurements**
BNP was measured quantitatively (pg/mL) using an ELISA assay (RayBiotech, USA). In this assay, a biotinylated BNP peptide is added into the samples. Plasma samples that contain endogenous BNP are subsequently added. The biotinylated BNP peptide competes with endogenous (unlabeled) BNP for binding to the anti- BNP antibody. After several washing steps, the remaining bound biotinylated BNP interacts with horseradish peroxidase (HRP)-streptavidin, which catalyzes a color development reaction measured as the extent of absorbance at 450 nm. In parallel, quantitative calibration is carried out using the provided BNP standards to generate a standard curve over the range of 0-1000 pg/mL BNP. Standard curve results are fit to a four-parameter logistic regression model. All plasma samples are diluted 2X to keep BNP levels within the dynamic range of the assay.

**Electrocardiogram**
EKGs were recorded in awake HFpEF and control mice using an EMKA tunnel system (Emka Technologies). Mice were injected (i.p.) with isoproterenol (5mg/kg body weight) and EKGs were recorded for 1 h. An arrhythmia score was assigned to each animal (0: no arrhythmias, 1: single PVCs, 2: coupled PVCs, 3: triple PVCs or non-sustained VT, 4: sustained VT) (2).

**Animals**
Male C57BL6/J mice were used in this study.

**Isolation of cardiac myocytes**
Cardiac myocytes were isolated as described previously(1, 3, 4). C57BL6/J mice were euthanized using isoflurane (5%) inhalation anesthesia via a vaporizer. 20 minutes prior to euthanasia mice were injected with an intraperitoneal heparin bolus (1000 U/kg). A thoracotomy was performed, and hearts were excised during deep anesthesia. Hearts were quickly immersed in ice-cold isolation buffer containing 130 mM NaCl, 5.4 mM KCl, 0.5 mM MgCl2, 0.33 mM $NaH_2PO_4$, 10 mM D-glucose, 10 mM taurine, 25 mM HEPES and 0.5 mM EGTA (pH 7.4, adjusted with NaOH). The aorta was canulated and hearts were then mounted on a Langendorff perfusion system. Hearts were perfused with isolation buffer containing 0.5 mM EGTA buffer for 5 minutes at 37ºC, before perfusion was switched to EGTA-free isolation buffer supplemented with 1 mg/mL collagenase (type II; Worthington Biochemical, USA), 0.06 mg/mL protease XIV, 0.06 mg/mL trypsin, and 0.3 mM $CaCl_2$. After 6-8 minutes of enzymatic perfusion the heart was dismounted. The ventricles were transferred to isolation buffer supplemented with 2 mg/ml BSA and 20 mM 2,3-butanedione monoxime and cut into small pieces. Mechanical dissociation with a fire polished Pasteur pipette was performed to achieve further dissolution of the ventricular tissue. The cell suspension was then filtered through a nylon mesh with a pore size of 300 µm. Ventricular myocytes were allowed to pellet by sedimentation, resuspended in NT solution, and were used within 4 hours of isolation. All procedures and protocols involving animal use were approved by the Institutional Animal Care and Use Committee of the University of Maryland School of Medicine.

**Measurements of $Ca^{2+}$ sparks and waves and stimulated $Ca^{2+}$ transients**
$Ca^{2+}$ sparks and waves were measured in freshly isolated ventricular cardiac myocytes from FM-HFpEF and age-matched C57BL6/J mice as described previously. Myocytes were loaded with the membrane-permeable fluorescent $Ca^{2+}$ indicator fluo4- AM (2-10 µmol/L) for 20 minutes at room temperature. After 20 minutes of de-esterification cells were seeded on a laminin coated coverslip and mounted on an inverted laser scanning confocal microscope (Nikon A1R). Cells were continuously perfused with modified Tyrode's solution (in mM: NaCl 130, KCl 5.4, $CaCl_2$ 1.8, $MgCl_2$ 0.5, $NaH_2PO_4$ 0.33, Glucose 5, HEPES 5, pH 7.4 adjusted with NaOH). Ventricular myocytes were electrically stimulated using platin electrodes in the bath connected to a pacing device (Myopacer, IonOptix). Steady state intracellular $Ca^{2+}$ handling was established by 2 minutes of

electrical field stimulation at a frequency of 1 Hz. $Ca^{2+}$ sparks and waves were elicited using the 488 (Argon) laser of the Nikon A1R in line scan mode. Spontaneous $Ca^{2+}$ sparks and waves were recorded for 30 s, data were collected and analyzed using Nikon elements software and the spark master(5) plugin for Fiji/image J (NIH). Stimulated $Ca^{2+}$ transients were recorded at 1 Hz and 4 Hz preceded by 1 minute of steady-state stimulation at either frequency.

### *Measurements of $[Ca^{2+}]_i$*

$[Ca^{2+}]_i$ in atrial and ventricular myocytes was measured as described previously(1, 3). Freshly isolated myocytes were loaded with the $Ca^{2+}$ indicator Fura2-AM (5-10 µM) for 20 minutes at room temperature. After 20 minutes of de-esterification cells were seeded on a laminin coated coverslip. Cells were mounted on an inverted microscope (Nikon Eclipse Ti2) connected to an EMCCD camera (Princeton Instruments Pro EM HS) and a rapid-switching illuminator with a 300 W xenon light source (DG5-plus, Sutter). Wide field imaging was achieved with 40X objective (Nikon S Fluor, Oil, 1.30 NA) using two excitation wavelengths (340 nm and 380 nm) by fast switching scanning mirrors and narrow bandwidth excitation filters (340 nm ± 10 nm; 380 nm ± 10 nm). Emission light was collected at 510 ± 40 nm. Data were acquired using Nikon NIS-Elements software. Fura2 fluorescence ($F_{340/380}$) was collected after steady-state $Ca^{2+}$ transients were established for 1 minute during external field stimulation. Cells were continuously perfused with modified Tyrode's solution (in mM: NaCl 130, KCl 5.4, $CaCl_2$ 1.8, $MgCl_2$ 0.5, $NaH_2PO_4$ 0.33, Glucose 5, HEPES 5, pH 7.4 adjusted with NaOH). $Ca^{2+}$ transients were elicited by external field stimulation (Myopacer, 2 ms, 20 V). After completion of measurements calibration of the $F_{340/380}$ signal was performed in situ in each cell. Myocytes were permeabilized with 2 µmol/L ionomycin and immobilized by cytochalasin D (50 µmol/L) and EGTA (1 mmol/L). Myocytes were perfused with a 0 and 10 mmol/L $Ca^{2+}$ modified Tyrode's solution (see above) containing ionomycin (2 µmol/L), cytochalasin D (50 µmol/L) and EGTA (1 mmol/L) to determine $F_{min}$ and $F_{max,}$ respectively. The $F_{340/380}$ signal from each cell was then calibrated using the following equation(6, 7):

$$[Ca^{2+}]_i: k_d * \beta * \frac{F - F_{min}}{F_{max} - F}$$

where $F_{min}$, $F_{max}$ and ß have their usual meaning. $k_d$ = 0.25 was obtained as described previously(8).

### Airy scan sub-diffraction super-resolution imaging

Imaging was carried out with a Zeiss LSM 880 confocal microscope equipped with an Airyscan super resolution imaging module using a 63/1.40 Plan-Apochromat Oil differential interference contrast M27 objective lens (Zeiss MicroImaging).

Freshly isolated ventricular myocytes were loaded with the membrane dye di-8 ANEPPS (5 μmol/L) as described previously. Cells from each animal were imaged within 1.5 h after cell isolation. For transverse-axial tubule (TAT) analysis, the NIH open-source Fiji platform was used(9). For each image of ventricular myocyte, the cellular interior region of interest (ROI) was selected using the polygon selection tool to exclude the cellular surface-membrane di-8 ANEPPS signal. Each obtained ROI was analyzed using a Fiji macro derived from(10, 11) and optimized for the signal-to-noise and resolution of the AiryScan images:

```
run("Add to Manager");
run("Enhance Contrast", "saturated=0.35");
run("Measure");
run("Duplicate…", " ");
run("Clear Outside");
run("Subtract Background…", "rolling=5");
run("8-bit");
run("Statistical Region Merging", "q=100 showaverages");
setThreshold(20, 255);
run("Convert to Mask");
run("Skeletonize (2D/3D)");
run("Directionality", "method=[Fourier components] nbins=180 histogram=-45
display_table"); run("Analyze Skeleton (2D/3D)");
```

Analysis outputs include Branch lengths, branch counts, junction counts, directionality histograms, ROI areas, and micron to pixel ratios were saved in excel files for each cell. These data in the excel files were parsed, analyzed, and plotted using Matlab (MathWorks, USA).

### *Tissue lysates and Western blot*

For tissue lysates, left ventricle obtained from C57BL/6 mice were cut into small pieces in ice-cold PBS supplemented with protease inhibitors (PBS-PI; cOmplete Mini EDTA-free; ROCHE). Samples were moved to 1.5 ml Eppendorf tubes suspended in PBS-PI and centrifuged at 20,000 x g at 4°C for 10 min. The pellets were resuspended in 2 x SDS sample buffer (Thermo Fisher Scientific) supplemented with 5% β-mercaptoethanol (Millipore) and homogenized using Eppendorf tube fitting pestles (SP SCIENCEWARE) until the suspension appeared homogenous. Samples were incubated at 100°C for 10 min and then centrifuged 20,000 x g at 4°C for 5 min. Supernatants were used for SDS PAGE. Protein concentrations were measured directly in the samples using a NanoDrop 1000 spectrophotometer (Thermo Fisher Scientific). 50 μg of protein per sample was separated on 4-20% gradient Novex Tris-glycine polyacrylamide gels (Thermo Fisher Scientific), and transferred onto polyvinylidene fluoride membranes (Bio-Rad Laboratories). Membranes were blocked in 5% blocking-grade nonfat dry milk (Bio-Rad Laboratories) in PBS-Tween20 and incubated with primary antibodies overnight at 4°C. NIR fluorescent secondary antibodies were used and Licor Revert total protein detection was employed. Membranes were imaged on a Licor

Clx imager (Li-Cor Biosciences, USA) with antibody fluorescence quantification, normalized to total protein, using Licor ImageStudio. Antibodies used for Western blotting were: anti-$Ca_v$1.2 (Alomone), anti-phospholamban (Invitrogen MA3-922), anti-phospho-phospholamban (Serine 16. Abcam AB15000), anti-serca2a (Invitrogen MA3-919), anti-calsequestrin (Invitrogen PA1-913), anti-ryanodine receptor type 2 (Alomone, ARR-002) and anti phospho- ryanodine receptor type 2 (Serine 2808, Invitrogen PA5-105712).
